# Concurrent representations of reinstated and transformed memories and their modulation by reward

**DOI:** 10.1101/2023.12.22.573008

**Authors:** Heidrun Schultz, Hanna Stoffregen, Aroma Dabas, María Alcobendas, Roland G. Benoit

**Affiliations:** Max Planck Institute for Human Cognitive and Brain Sciences, Leipzig, Germany; Charité-Universitätsmedizin Berlin, Berlin, Germany; University of Colorado Boulder, USA

**Author notes:** These authors contributed equally to this work. **Corresponding authors**: Heidrun Schultz, Max Planck Institute for Human Cognitive and Brain Sciences, Stephanstr. 1A, 04103 Leipzig, Germany. Roland G. Benoit, University of Colorado Boulder, Department of Psychology and Neuroscience & Institute of Cognitive Science, UCB 344, MUEN PSYCH Building D418, University of Colorado, Boulder, Colorado 80309.

**Keywords:** reinstatement, memory transformation, memory generalization, medial prefrontal cortex, medial temporal lobe

## Abstract

An integral part of episodic retrieval is the reinstatement of neural activity that was present in the medial temporal lobe during encoding. However, neural memory representations do not remain static. Consolidation promotes the transformation of representations that are specific to individual episodes towards more generalized representations that reflect commonalities across episodes. Moreover, reward has been shown to augment episodic memory by enhancing consolidation, and it may accelerate the transformation of neural memory representations. We investigated this account with n=40 human participants using fMRI and an associative memory task. They encoded pictures of objects, each with one of four recurring scenes. Two scenes led to high reward, two led to low reward. The next day, participants encountered the objects again and retrieved the scenes from memory. Using representational similarity analysis, we demonstrate that retrieval is concurrently accompanied by the reinstatement of original neural representations and the activation of transformed, more generalized memories. Specifically, the parahippocampal cortex reinstates scene-specific patterns from the encoding phase during successful retrieval. In contrast, activity patterns in the medial prefrontal cortex and anterior hippocampus reflect transformed memories: They become more similar to each other for memories sharing the same scene, independent of memory success. Importantly, high reward enhances memory transformation in the anterior hippocampus. The brain thus maintains complementary memory representations: An episodic representation that resembles the original encoding pattern, and a generalized representation that summarizes commonalities across memories - in part for particularly valuable information.

## Introduction

The human ability to retain memories is remarkable: Seemingly effortlessly, we can recall the picnic we had yesterday in rich detail. At the same time, we also know how picnics generally work, allowing us to easily plan for an upcoming event that may take place on the next weekend. This is because, over time, the commonalities across similar episodes are extracted and the memories are thus transformed into more generalized knowledge. Such knowledge can take the form of, for example, mental schemas, scripts, or categories (Ghosh & Gilboa, 2014; Gilboa & Marlatte, 2017).

Distinct forms of memories are reflected in distinct neural patterns. On the one hand, episodic recall is accompanied by reinstatement of the original encoding activity. This has especially been shown in content-sensitive regions of the medial temporal lobe (MTL), including the parahippocampal cortex (PHC) (Schultz et al., 2019; Schultz, Sommer, et al., 2022; Staresina et al., 2012) and the (posterior) hippocampus (HC) (Bone & Buchsbaum, 2021).

On the other hand, neural patterns associated with generalized memories do not reflect the encoding activity of any individual episode. Over time, memories that share common features undergo a transformation so that their neural representations become more similar. This has been shown in the medial prefrontal cortex (mPFC) and HC (Audrain & McAndrews, 2022; Tompary & Davachi, 2017).

The mPFC may thus represent transformed memories in the form of generalized knowledge structures (Ghosh & Gilboa, 2014; Gilboa & Marlatte, 2017; Milivojevic et al., 2015; Paulus et al., 2021), whereas the hippocampus may contain both types of memory representations. Notably, there is some evidence for a functional specialization within the hippocampus – with more general memories, such as an episode’s gist, being supported by the anterior HC, and more detailed episodic memories being more reliant on the posterior HC (Collin et al., 2015; Gilboa & Moscovitch, 2021; Guo & Yang, 2020; Poppenk et al., 2013; Sekeres et al., 2018).

Tompary and Davachi (2017) recently investigated memory transformation through multivariate pattern analysis of fMRI data. They had participants encode pairs of unique objects with one of four recurring scenes. Either immediately following encoding or one week later, participants were cued with the objects to retrieve the scenes from memory. In both the mPFC and MTL, neural patterns during retrieval were more similar to each other for objects that had shared the same scene (retrieval-retrieval similarity). Critically, this was only the case during the delayed memory test, indicating that the memories underwent a transformation over time. The authors concluded that consolidation promotes representational convergence of memories that share overlapping features, which may be an important building block for memory generalization.

We here seek to build on this work to address three questions. First, the reported increase in retrieval-retrieval similarity for objects sharing the same scene (Tompary & Davachi, 2017) may not necessarily reflect memory generalization. Instead, it may be a byproduct of scene reinstatement, i.e. the reactivation of the same scene-specific encoding pattern during retrieval (Mack & Preston, 2016; Wing et al., 2015): If the same scene encoding pattern is reinstated in two retrieval trials, these trials may then be more similar to each other, thus potentially driving retrieval-retrieval similarity. Such an effect could even increase over time, given the mPFC’s time-dependent role in memory retrieval (Barry et al., 2018; Bonnici et al., 2012; Bonnici & Maguire, 2018; Sekeres et al., 2018; Sommer, 2017). Here, we address this issue by also examining scene reinstatement as a potential alternative account for the effect reflected in retrieval-retrieval similarity.

Second, we investigate whether generalized memory representations can be expressed even in the absence of successful retrieval of individual episodes. Given that generalization entails loss of episodic detail (Sekeres et al., 2018), we suggest that such generalized representations may be activated by an episodic retrieval cue, even if these representations do not provide sufficient detail to drive successful episodic retrieval.

Third, previous work has been agnostic with regards to the drivers of memory generalization. We here suggest that memory transformation is promoted by reward. Neural replay during consolidation appears to be critical to generalization (Kumaran et al., 2016; Liu et al., 2019), and rewarded memoranda are preferably replayed post encoding (Gruber et al., 2016; Sterpenich et al., 2021). In general, we thus suggest that reward may facilitate memory generalization.

Specifically, the mPFC has been implicated not only in representing generalized knowledge structures (Gilboa & Marlatte, 2017) but also in reward processing (Haber & Knutson, 2010). Indeed there is evidence that mPFC representations may be shaped by value (Baram et al., 2021; Moneta et al., 2023; Paulus et al., 2021). Similarly, the anterior HC may not only be particularly involved in representations of broad, general memories, but may also process motivationally relevant aspects of a memory - such as reward (Poppenk et al., 2013). We thus hypothesize that reward facilitates the representational convergence of memories that share overlapping features.

To address these questions, we conducted a two-day fMRI study with n=40 participants. On day one, participants engaged in an incidental encoding task (adapted from Gruber et al., 2016), in which they associated a series of single objects with one of four recurring scenes. Two of the scenes led to high reward, and the two other scenes to low reward. The next day, participants returned for a surprise scene recall task (adapted from Tompary & Davachi, 2017). Here, participants were cued with each object to recall the associated scene.

First, we tested the hypothesis that scene-specific reinstatement is present in the PHC and posterior HC, but not the mPFC. Successful retrieval should thus be associated with greater encoding-retrieval similarity for trials that share the same scenes, compared to those that share different scenes. Second, we tested the hypothesis that the mPFC and anterior HC, but not the PHC, represent transformed memories. This would be reflected in increased retrieval-retrieval similarity for objects that had shared the same encoding scene. Third, we expected this effect to be greater for overlapping memories that had been highly rewarded.

## Method

### Sample

A total of n=42 volunteers took part in the study. Of these, two were excluded from data analysis (one due to equipment failure, one did not return for the second day). We thus report data from n=40 participants (mean age: 26.25 years, age range: 19-35 years, 28 women, 12 men). They were native German speakers with normal or corrected-to normal vision and without a history of psychiatric or neurological disorder. The study protocol was approved by the ethics committee of the medical faculty of the University of Leipzig (171/19-ek), and all participants provided written informed consent prior to participating. They received 9€/hour and an additional bonus of up to 15€, depending on their performance during the encoding task.

### Procedure and tasks

Participants took part in two fMRI sessions on consecutive days (mean delay between fMRI sessions: 22h 35min, range: 19h 30min – 26h 0min). On day one, they engaged in the incidental encoding task; on day two, they returned for the surprise scene recall task.

#### Day 1: Incidental encoding task

The incidental encoding task (Figure 1A) was adapted from Gruber et al. (2016) (see also Schultz et al., 2023). In each trial, participants viewed one of 160 objects paired with one of four recurring scenes: A circus, a basketball court, an office, and a swimming pool. Participants were asked to mentally simulate a scene-specific action and respond to a corresponding question (e.g. “Would the object float on water?” for the swimming pool scene). Correct responses yielded a high or low reward. Importantly, two of the scenes were always paired with high reward (2 points), while the other two scenes were always paired with low reward (0.02 points). Points were later converted to monetary reward. Allocation of scenes to reward magnitudes were instructed prior to scanning, and counterbalanced across participants. Each object-scene pair was presented twice, with the repetition occurring within the same run. This yielded a total of 320 trials over four runs.

**Figure 1.**
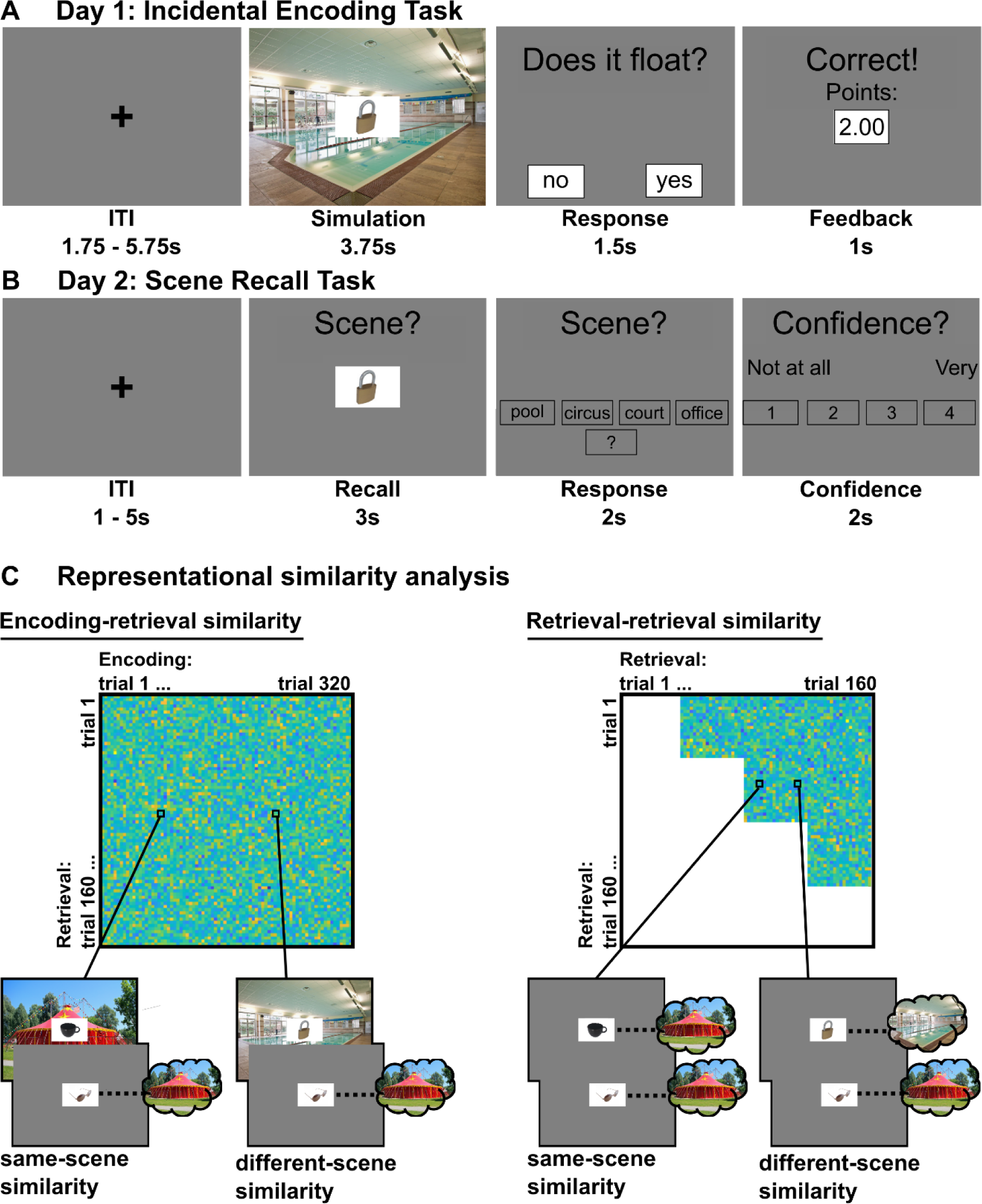
**A**. Example trial for the incidental encoding task. After a variable inter-trial interval of 1.75 – 5.75s, participants viewed one of 160 objects on top of one of four scenes, and mentally engaged in a scene-specific simulation task (here: “Would the object float on the water?”) for 3.75s. They had 1.5s to respond “yes” or “no”, and received reward feedback (high: 2.00 points, low: 0.02 points) for 1s for each correct response. **B**. Example trial for the scene recall task. After a variable ITI, participants viewed one of the objects from the encoding phase for 3s and tried to retrieve the scene from memory. They then had 2s to respond by choosing a label presented on the screen (e.g. “swimming pool”). If they chose a scene label, they then had 2s to rate the confidence of their choice on a scale of one (“not at all confident”) to four (“very confident”). **C**. Representational similarity analysis. We used two measures to assess similarity between trials that either shared a scene (same-scene similarity) or did not (different-scene similarity). To test reinstatement of scene-specific activity from the encoding phase, we assessed encoding-retrieval similarity. That is, we correlated pairs of retrieval trials with encoding trials that shared the same scene or not. To test for memory generalization based on overlapping features (i.e. the same scene), we assessed retrieval-retrieval similarity. That is, we correlated pairs of trials from the retrieval phase that had either shared a scene (same-scene similarity) or had not (different-scene similarity). For copyright reasons, we here display photographs that are similar to the actual experimental stimuli (scene images from http://pixabay.com; object images by the investigators).

#### Day 2: Scene recall task

The scene recall task (Figure 2A) was adapted from Tompary and Davachi (2017) (see also Schultz et al., 2023). In each trial, one of the objects from the encoding task was presented, and participants were asked to recall the scene that it had been paired with. They responded with one of five choices (verbal labels for the four scenes plus a “don’t know” option). If they responded with a scene, they were also asked to rate the confidence of their choice on a scale from one (not at all) to four (very). Each object was presented once, yielding a total of 160 trials across four runs.

**Figure 2.**
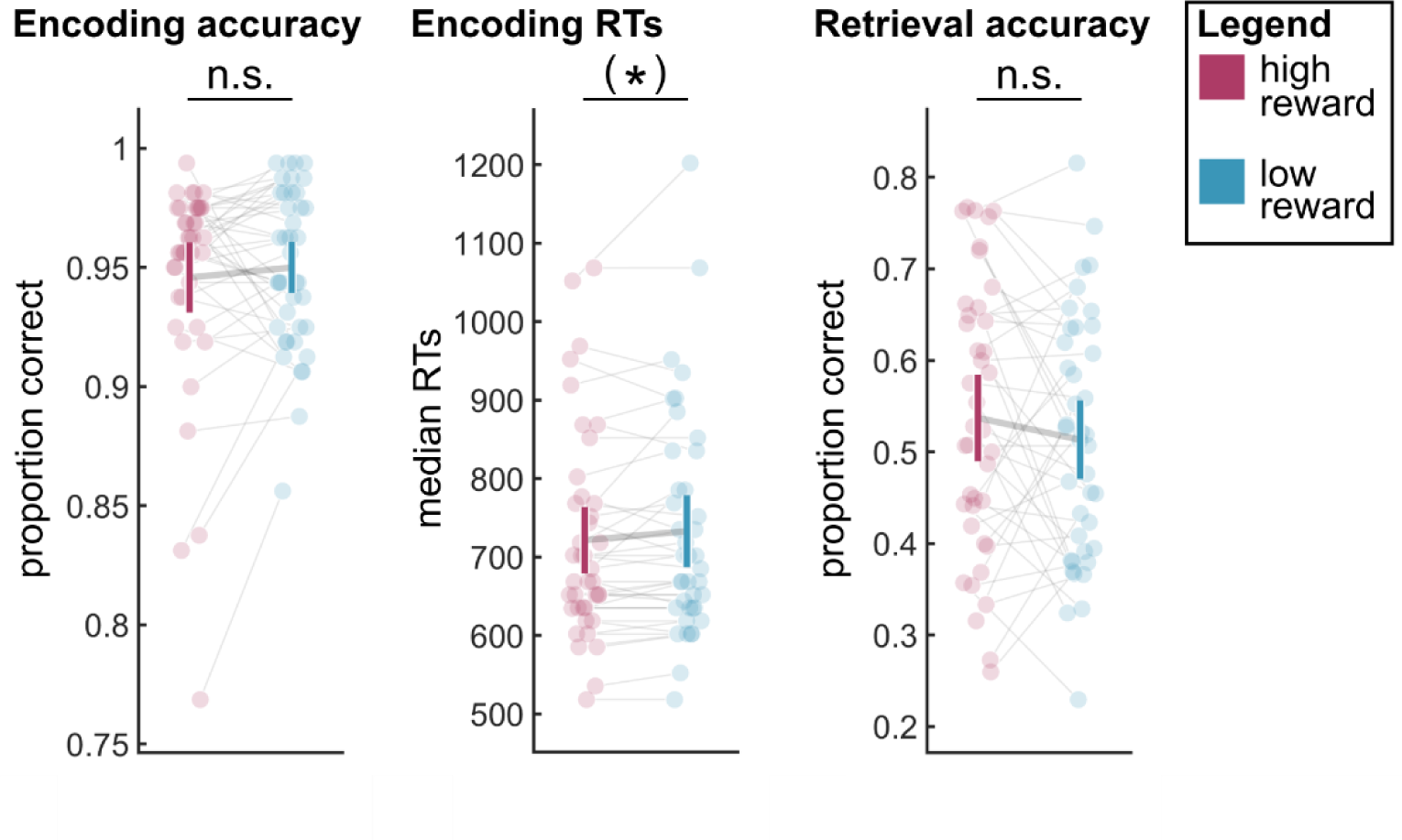
Behavioral results. Left panel: Accuracy during the incidental encoding task. Middle panel: Median response times (RTs) for correct responses during the incidental encoding task. Right panel: Accuracy during the scene recall task. Error bars indicate 95% confidence interval. Abbreviations: n.s., not significant; (*) .05<*p*<.1.

Day two also included an unscanned recall task for the reward magnitude (Schultz et al., 2023), as well as questionnaires. These are not part of the present report.

### Implementation

Tasks were implemented in Octave (RRID:SCR_014398) and the Psychophysics Toolbox (RRID:SCR_002881). We note that, due to a bug in the code, object images in both the incidental encoding task and scene recall task were stretched to an approximately 4:3 format.

### Behavioral analysis

For the incidental encoding task, we computed proportions of correct responses as well as median response times for correct responses, separately for high and low reward trials. For the scene recall task, we computed proportions of correct scene recall, separately for high and low reward trials. All further analyses included only trials for which the encoding task had been answered correctly on both repetitions. We tested for effects of reward using paired *t*-tests on all three behavioral measures. Statistical analyses were conducted in R (RRID:SCR_001905) and RStudio (RRID:SCR_000432).

### MRI acquisition

MRI data were acquired on a Siemens Prisma 3T system. Functional data were scanned using a whole-brain T2*-weighted gradient-echo, echo-planar pulse sequence (2mm isotropic voxels, 72 interleaved slices, TR=2000ms, TE=25ms, multiband acceleration factor=3). On each day, five functional runs were acquired: One run of 120 volumes of rest followed by either four runs of 526 volumes (task plus rest, day one) or 206 volumes (task, day 2). Gradient-echo fieldmaps were acquired at the beginning of each session. On day one, we also acquired a T1-weighted structural image (MPRAGE, 1mm isotropic voxels). Additional structural scans (DWI, MP2RAGE) were acquired that were not analyzed for the present report.

### MRI preprocessing and first-level statistics

The MRI data were first converted to the Brain Imaging Data Structure (BIDS) (Gorgolewski et al., 2016). Preprocessing was performed using fMRIPrep 21.0.2 (Esteban et al., 2019; Esteban, Markiewicz, Goncalves, et al., 2022) (RRID:SCR_016216) based on Nipype 1.6.1 (Esteban, Markiewicz, Burns, et al., 2022; Gorgolewski et al., 2011) (RRID:SCR_002502).

The T1-weighted image (T1w) was corrected for intensity non-uniformity, skull-stripped, and segmented into gray matter (GM), white matter (WM), and cerebrospinal fluid (CSF). It was then normalized to standard space (MNI152NLin2009cAsym). From the functional data, first, a reference image was estimated for use in the motion correction and co-registration steps. The functional data were slice-time corrected to the middle temporal slice, motion-corrected, and corrected for susceptibility distortions using the fieldmap acquired at the start of each session. Functional data were then co-registered to the T1w using boundary-based registration (Greve & Fischl, 2009) with six degrees of freedom (for further preprocessing details, see https://fmriprep.org/en/21.0.2/).

We conducted the further processing in MATLAB (RRID:SCR_001622) and SPM12 (RRID:SCR_007037). Specifically, we set up two sets of first-level general linear models (GLMs) on the unsmoothed, non-normalized data: one set for the encoding and one for the retrieval data. Each trial was estimated in a separate GLM (Mumford et al., 2012), with a single regressor on the simulation onset respectively recall onset. Each model also contained categorical regressors for all other onsets, separately for each of the conditions (HR: high reward/remembered, HF: high reward/forgotten, LR: low reward/remembered, LF: low-reward/forgotten) as well as a categorical regressor encompassing all button presses. All of these regressors were convolved with the hemodynamic response function (HRF). Additionally, each model included a set of seven non-convolved noise regressors extracted during preprocessing, i.e. the six rigid motion regressors (three translations, three rotations) as well as framewise displacement. Functional runs were concatenated, and session constants were included in the models. The resulting beta maps for each trial of the encoding and retrieval sessions were converted to *t* maps. Finally, the *t* maps were minimally smoothed with a Gaussian kernel of 2mm full width at half maximum (Dimsdale-Zucker & Ranganath, 2018).

### Regions of interest

We employed bilateral anatomical masks of the PHC, the whole HC as well as its anterior and posterior subdivisions, and mPFC. For the PHC, anterior HC, and posterior HC masks, we automatically segmented each participant’s T1w using ASHS (Yushkevich et al., 2015) and the Penn Memory Center 3T ASHS Atlas for T1-weighted MRI (Xie et al., 2016). We reviewed each individual segmentations and found that the automated process led to a successful outcome for our participant population.

The PHC masks were then manually adjusted according to guidelines suggested by (Pruessner et al., 2002). The anterior and posterior HC masks were also combined into a single HC mask. All masks were then resampled to each participant’s functional space. For visualization (Figure 3, 4A-B), the masks were warped into standard space using the transformation matrix from the T1w normalization, and averaged across participants.

**Figure 3.**
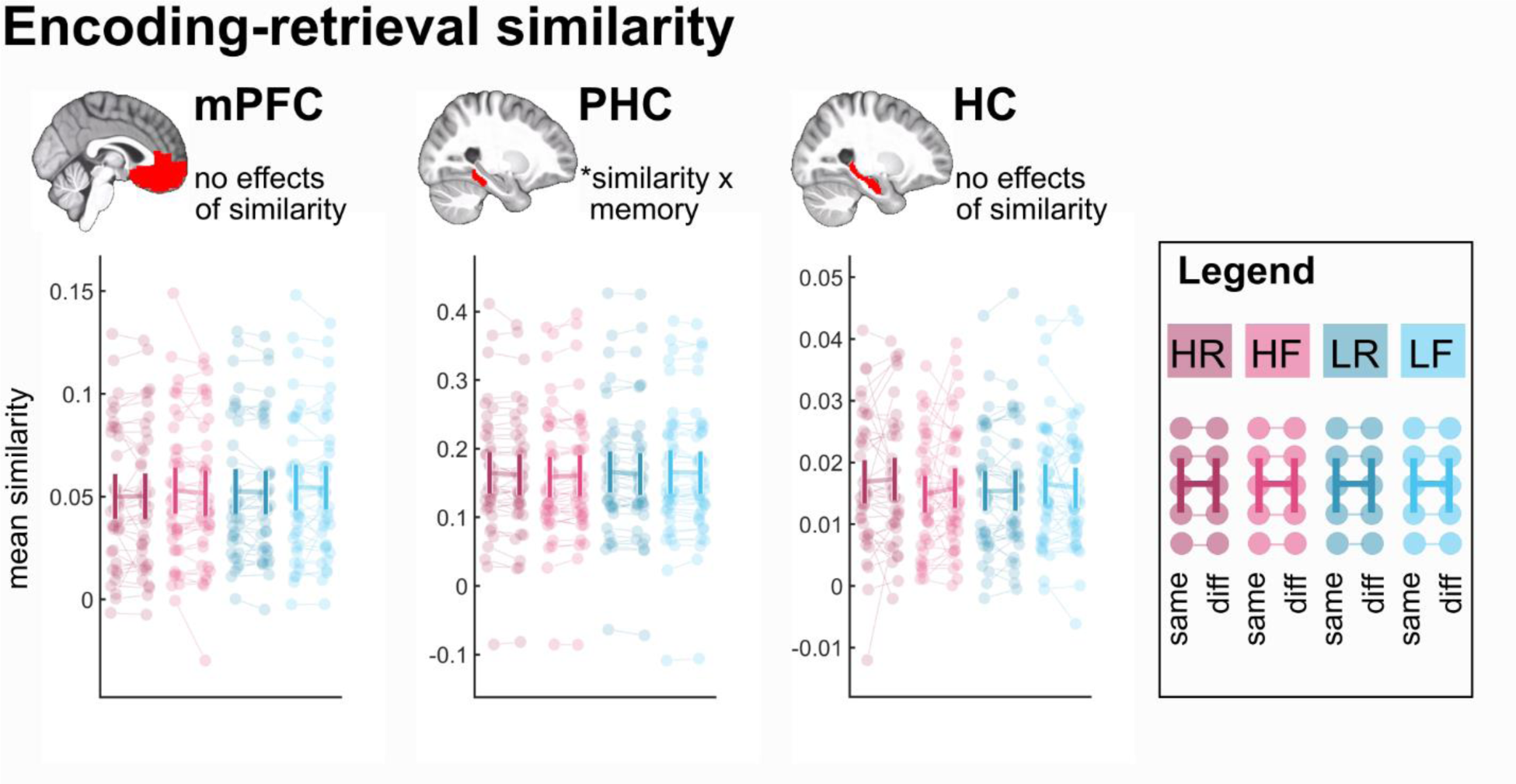
Encoding-retrieval similarity. Average encoding-retrieval similarity values in the three a-priori ROIs: mPFC, PHC, and HC. Notes refer to effects from three-way repeated-measures ANOVAs with the factors reward, memory, and similarity (see main text for details). Error bars indicate 95% confidence interval. Abbreviations: HR: high-reward/remembered; HF: high-reward/forgotten; LR: low-reward/remembered, LF: low-reward/forgotten; same: same-scene similarity, diff: different-scene similarity; \**p*<.05.

**Figure 4.**
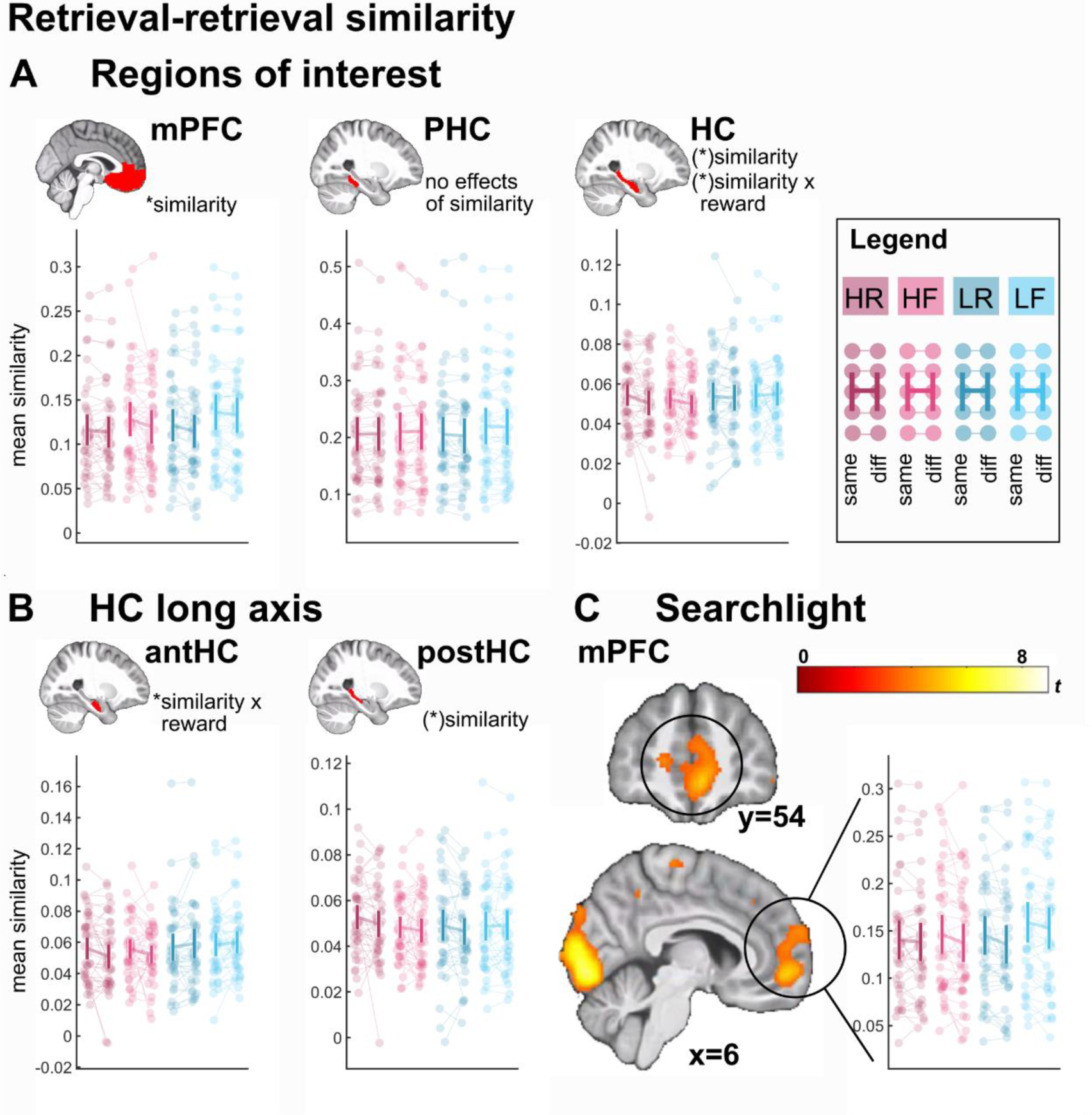
Retrieval-retrieval similarity. **A**. Average retrieval-retrieval similarity values in the three a-priori ROIs: mPFC, PHC, and HC. **B**. Average retrieval-retrieval similarity in the anterior vs. posterior portion of the HC. **C**. Results of whole-brain searchlight analysis for the main effect of similarity (same-scene similarity > different-scene similarity across levels of reward and memory). Average retrieval-retrieval similarity was extracted for visualization. Display threshold *p*<.001 unc., *k*=5 voxels. Notes refer to effects from three-way repeated-measures ANOVAs with the factors reward, memory, and similarity (see main text for details). Error bars indicate 95% confidence interval. Abbreviations: HR: high-reward/remembered; HF: high-reward/forgotten; LR: low-reward/remembered, LF: low-reward/forgotten; same: same-scene similarity, diff: different-scene similarity; \**p*<.05, (*) .05<*p*<.1.

For the mPFC mask, we used the Brainnetome atlas (Fan et al., 2016) and combined left and right medial area 11, 13, and 14 as well as left and right subgenual area 32. This mask was warped from standard space into each participant’s single subject space using the inverse transformation from the T1w normalization and the functional reference image (see above).

### Representational similarity analysis

From the minimally smoothed single-trial *t* maps, we extracted and vectorized *t* values from each ROI for each trial. Similarity was operationalized as the average Fisher-z-transformed Pearson correlation coefficient between pairs of trials. Our approach is illustrated in Figure 1C. For the MTL ROIs (PHC, HC), similarity values were calculated separately for the left and right hemisphere and then averaged.

First, to test for episodic reinstatement of scene-specific information, we computed *encoding-retrieval-similarity*. Here, we computed the average similarity between pairs consisting of an encoding and a retrieval trial, separately for the following conditions: High-reward remembered trial pairs sharing the same scene (HR-same) or different scene (HR-diff), and similarly for high-reward forgotten trial pairs (HF-same, HF-diff), low-reward remembered trial pairs (LR-same, LR-diff), and low-reward forgotten trial pairs (LF-same, LF-diff). We excluded trial pairs containing the same object as well as trials with incorrect encoding responses.

Second, to test for memory generalization, we computed *retrieval-retrieval similarity.* Here, we computed the average similarity between pairs of retrieval trials, separately for the following conditions: High-reward remembered trial pairs sharing either the same scene (HR-same) or different scenes (HR-diff), and similarly for high-reward forgotten trials (HF-same, HF-diff), low-reward remembered trials (LR-same, LR-diff), and low-reward forgotten trials (LF-same, LF-diff). We excluded trial pairs from the same fMRI run as well as trials with incorrect encoding responses.

Statistical analyses of the similarity values were conducted in R. To test for effects of our experimental manipulations on encoding-retrieval similarity and retrieval-retrieval similarity in each ROI, we submitted the mean similarity scores for each participant and condition to repeated-measures ANOVAs (R: afex::aov_ez) (Singmann et al., 2021) with the factors reward (high, low), memory (remembered, forgotten), and similarity (same scene, different scene). We conducted follow-up paired comparisons as necessary.

### Representational similarity analysis – complementary searchlight analyses

We complemented the above ROI analyses of encoding-retrieval similarity and retrieval-retrieval similarity with a searchlight analysis. To this end, we repeated the above analysis within a moving searchlight (3 voxel radius) centered on every voxel inside each participant’s brain mask. For each condition (i.e. HR-same, HR-diff, HF-same, HF-diff, LR-same, LR-diff, LF-same, LF-diff), the resulting fisher-z transformed correlation coefficients were written out as a statistical image. For each participant, we then combined these statistical images into first level contrast images: Following up on the ROI results, for encoding-retrieval similarity, we computed the interaction effect of memory and similarity ([1 -1 -1 1 1 -1 -1 1]). For retrieval-retrieval similarity, we computed the main effect of similarity ([1 -1 1 - 1 1 -1 1 -1]). These contrast images were normalized to standard space using the transformation matrix from the T1w normalization, and smoothed with a Gaussian kernel of 6mm full width at half maximum (SPM12). The resulting maps were then submitted to a second-level one-sample *t*-test in SPM12. Multiple comparisons correction was achieved through peak-level family-wise error correction (FWE) within the anatomical masks of the HC and PHC (encoding-retrieval similarity) and the mPFC mask (retrieval-retrieval similarity). Finally, to explore effects outside our ROIs, we also applied FWE correction across the whole brain.

## Results

### Behavioral results

Accuracy on the incidental encoding task was, as intended, near ceiling (Figure 2), and did not differ for high versus low reward trials (*t*_(39)_=0.766, *p*=.449). There was a trend for correct responses to be faster in the high than the low reward condition (*t*_(39)_=1.755, *p*=.087, Figure 2).

Participants correctly recalled around half of the scenes. While accuracy was numerically higher for high-reward trials (high reward: 53.7%, low reward: 51.3%, Figure 2), the difference was not significant (*t*_(39)_=1.147, *p*=.258).

### Encoding-retrieval similarity: The PHC reinstates scene-specific patterns during retrieval

We tested whether, during retrieval, the three ROIs would reinstate scene-specific patterns from the encoding phase. To this end, we calculated the average similarities between pairs of retrieval trials and encoding trials that either shared the same scene or not (same-scene versus different-scene similarity). Reinstatement would be reflected in greater same-scene than different-scene similarity. We expected this similarity effect for remembered trials, particularly in the PHC and HC, with the mPFC serving as control.

#### ROI analyses

In each of the three ROIs (mPFC, PHC, and HC), we computed a three-way repeated-measures ANOVA with the factors reward (high, low), memory (remembered, forgotten), and similarity (same scene, different scene). We report main effects of similarity as well as interactions that include the similarity factor, as other effects (such as e.g. a main effect of memory) do not reflect reinstatement of the specific scene.

The mPFC (Figure 3, left panel) did not show an effect involving the similarity factor (all *F*_(1,39)_≤0.482, all *p*≥.492).

The PHC (Figure 3, middle panel) showed an interaction of similarity and memory (*F*_(1,39)_=5.370, *p*=.026), reflecting a greater similarity effect for remembered than forgotten trials. No other effects involving the similarity factor were significant (all *F*_(1,39)_≤2.079, all *p*≥.157). To follow up on this interaction, we averaged over the reward factor, and compared same-vs. different scene similarity separately for remembered and forgotten trials. Same-scene similarity was significantly greater than different-scene similarity for remembered (*t*_(39)_=2.796, *p*=.008) but not forgotten trials (*t*_(39)_=0.617, *p*=.541). This pattern is consistent with reinstatement of scene-specific encoding activity during successful memory retrieval.

Overall, the HC (Figure 3, right panel) did not show any effects involving the similarity factor (all *F*_(1,39)_≤1.235, all *p*≥.273). However, previous research suggests that the posterior HC may be particularly involved in representing episodic detail (Poppenk et al., 2013). We therefore repeated the above analysis separately for the anterior and posterior HC. The anterior HC did not show any effects involving the similarity factor (all *F*_(1,39)_≤0.367, all *p*≥.548), while the posterior HC only showed a trend-level interaction of reward and similarity (*F*_(1,39)_=3.534, *p*=.068, all other *F*_(1,39)_≤0.891, all *p*≥.351, numerically greater similarity effect for low-reward than high-reward trials).

#### Complementary searchlight analysis

To follow up on our findings of scene-specific reinstatement during remembered trials in the PHC, we repeated the same analysis using a searchlight approach. The contrast for the interaction effect of memory and similarity (i.e. [1 -1 -1 1 1 -1 -1 1] on the conditions HR-same, HR-diff, HF-same, HF-diff, LR-same, LR-diff, LF-same, LF-diff) was computed on the single-subject level and submitted to a one-sample *t* test on the group level. However, at *p*<.001 uncorrected, this analysis yielded no significant voxels within a combined mask of the PHC and HC. Furthermore, no voxels survived FWE correction across the whole brain. We note that, due to the high interindividual variability in MTL anatomy (Pruessner et al., 2002), analyses in group space are typically less sensitive than analyses within individual MTL ROIs.

### Retrieval-retrieval similarity: The mPFC and anterior HC represent transformed memories

Next, we tested whether memory representations with overlapping features (i.e. the same scene) showed evidence for generalization. To this end, we calculated the average similarities between pairs of retrieval trials that either shared the same scene (same-scene similarity) or not (different-scene similarity). Memory generalization would be reflected in greater same-scene than different-scene similarity. We expect this similarity effect predominantly in the mPFC and HC, with the PHC serving as control.

#### ROI analyses

As with the encoding-retrieval similarity analysis above, we computed, for each of the ROIs, a three-way repeated-measures ANOVA with the factors reward (high, low), memory (remembered, forgotten), and similarity (same scene, different scene). We report main effects of similarity as well as interactions that include the similarity factor, as other effects (such as a main effect of memory) do not reflect generalization.

The mPFC (see Figure 4A) showed a significant main effect of similarity (same-scene similarity > different-scene similarity, *F*_(1,39)_=13.455, *p*<.001). This is consistent with the emergence of generalized memory representations for episodes that share overlapping features. No other effect involving the similarity factor was significant (all *F*_(1,39)_≤1.932, all *p*≥.172).

The PHC (see Figure 4A) did not yield an effect involving the similarity factor (all *F*_(1,39)_≤0.774, all *p*≥.384).

Overall, for the HC (see Figure 4A), we observed a trend-level main effect of similarity (same-scene similarity > different-scene similarity, *F*_(1,39)_=3.670, *p*=.063), qualified by a trend-level interaction of reward and similarity (*F*_(1,39)_=2.891, *p*=.097). This pattern reflected a larger similarity effect for high-reward than low-reward trials. Given that the anterior portion of the HC may be particularly involved in processing generalized information as well as reward (Guo & Yang, 2020; Poppenk et al., 2013), we repeated the above analyses in anterior vs. posterior portions of the HC (Figure 4B).

The anterior HC showed a significant interaction of reward and similarity (*F*_(1,39)_=9.743, *p*=.003). This is consistent with reward-enhanced generalization of overlapping memories. No other effect that included the similarity factor was significant (all *F*_(1,39)_≤1.213, all *p*≥.278). To follow up on this interaction, we averaged over the memory factor, and compared same-scene vs. different scene similarity, separately for high-reward and low-reward trials. Same-scene similarity was significantly greater than different-scene similarity for high-reward trials (*t*_(39)_=2.686, *p*=.011), but not for low-reward trials (*t*_(39)_=1.624, *p*=.112). The anterior HC thus showed a result pattern that was more pronounced than the result pattern in the whole HC ROI.

The posterior hippocampus, on the other hand, showed a trend-level main effect of similarity (same-scene similarity > different-scene similarity, *F*_(1,39)_=3.880, *p*=.056). No other effect including the similarity factor was significant ((all *F*_(1,39)_≤1.652, all *p*≥.206).

#### Complementary searchlight analysis

To corroborate our main findings of overall memory generalization in the mPFC, we conducted the same analysis using a searchlight approach. The contrast for the main effect of similarity (i.e. [1 -1 1 - 1 1 -1 1 -1] on the conditions HR-same, HR-diff, HF-same, HF-diff, LR-same, LR-diff, LF-same, LF-diff) was computed on the single-subject level and submitted to a one-sample t-test on the group level. We applied small-volume correction across the anatomical mPFC mask. This analysis yielded a significant peak within the mPFC (MNI coordinates: [6 54 -6], *t*_(39)_=5.212, *p*_SVC_=.003, Figure 4C). We visualized this effect by extracting the mean similarity values across the entire cluster (thresholded at *p*<.001) for each condition and subject. The result pattern resembles the one reported for the anatomical mPFC ROI (see above). In addition, two further peaks survived FWE-correction across the whole brain: The left postcentral gyrus ([-44 -22 54, *t*_(39)_=8.890, *p*_FWE_<=.001), and the bilateral occipital cortex ([0 -88 2], *t*_(39)_=8.325, *p*_FWE_<=.001).

## Discussion

With the present fMRI study, we set out to examine whether the brain concurrently represents episodic memory representations as well as more generalized memory representations that encode commonalities across episodes.

First, we examined the reinstatement of scene patterns from the encoding experience, thought to reflect episodic memory. We observed evidence for memory reinstatement in the PHC, but not the mPFC or HC. Second, we tested for memory transformation – i.e. the representational convergence of neural retrieval patterns for memories that share overlapping features. This is thought to reflect the shift from individual episodes into generalized memory. We found evidence for such memory transformation in the mPFC and HC, but not the PHC. Intriguingly, memory transformation in the mPFC affected all memories, regardless of reward magnitude or retrieval success. In contrast, memory transformation in the anterior HC was enhanced by reward. We note, however, that unlike previous studies (e.g.) (Gruber et al., 2016; Schultz, Yoo, et al., 2022; Wittmann et al., 2005) we did not observe a reward effect on behavioral retrieval accuracy, and only a marginal effect on response times during encoding (see also Schultz et al., 2023).

Our results thus extend our knowledge derived from studies that had tested retrieval after a consolidation period of three days (Audrain & McAndrews, 2022) and one week (Tompary & Davachi, 2017). Here, we demonstrate (i) that memory transformation has already taken place after one day, (ii) that it is partly enhanced by reward, and (iii) that generalized memory representations can be activated even in absence of successful retrieval of a particular episodic memory.

Importantly, did our analysis of retrieval-retrieval similarity truly reflect transformed memories? Transformation implies two things: That a representation has changed over time, and that it is dissimilar to the original encoding pattern. Previous studies have focused on the first implication and demonstrated that memories sharing the same scene become more similar to each other over time (Audrain & McAndrews, 2022; Tompary & Davachi, 2017). The second implication is equally critical. This is because retrieval-retrieval similarity could also be a consequence of common scene reinstatement: If the same-scene-specific encoding pattern is reinstated in two retrieval trials, these would also be similar to each other. Such an effect would not reflect a transformation away from the original encoding pattern.

To address this, we analyzed concurrent scene reinstatement (i.e. scene-specific encoding-retrieval similarity). Our results imply that retrieval-retrieval similarity was not merely driven by scene reinstatement: First, there was little topographical overlap between the two effects, with scene reinstatement predominantly in the PHC, and retrieval-retrieval similarity predominantly in the mPFC and (anterior) HC. Second, scene reinstatement was modulated by memory success whereas retrieval-retrieval similarity was not. This pattern indicates that our findings in the mPFC and anterior HC truly reflect a transformed memory representation rather than a reinstatement of a shared encoding pattern.

Memory transformation – in the sense of representational convergence of overlapping memories - has previously been observed for the mPFC (Audrain & McAndrews, 2022; Tompary & Davachi, 2017). These findings are consistent with a role of the mPFC in representing generalized knowledge structures (Ghosh & Gilboa, 2014; Gilboa & Marlatte, 2017; Paulus et al., 2021).

Generalization may be driven by replay of episodic memories during consolidation. This process would allow the cortex to extract commonalities across similar memories and to store these as more generalized representations (Kumaran et al., 2016; Liu et al., 2019; Sekeres et al., 2018). Here, we show that such effects do not require three (Audrain & McAndrews, 2022) or seven days (Tompary & Davachi, 2017) to emerge. Instead, they are already present a single day after encoding.

Notably, in the mPFC, the activation of transformed memory representations was independent of memory success. This is somewhat at odds with previous studies that reported memory transformation for correctly recalled trials only (Audrain & McAndrews, 2022; Tompary & Davachi, 2017). However, both of these studies included control analyses for trials without overt scene recall (recognition trials, Tompary & Davachi, 2017; forgotten trials, Audrain & McAndrews, 2022). In both cases, the mPFC patterns were numerically consistent with representational convergence, though they were statistically inconclusive. If the mPFC encodes representations that generalize across episodes that share common content (Audrain & McAndrews, 2022; Gilboa & Marlatte, 2017; Sekeres et al., 2018), these would get activated whenever one of these episodes is being probed. However, given that these representations abstract away from unique features that are specific to individual episodes, they would not contain sufficient episodic detail to drive episodic recall.

Contrary to our hypothesis, reward did not foster memory transformation in the mPFC, but only in the anterior HC. Reward has been shown to increase neural replay (Gruber et al., 2016; Sterpenich et al., 2021) and promote consolidation (Murayama & Kitagami, 2014; Murayama & Kuhbandner, 2011; Spaniol et al., 2014; Wittmann et al., 2005). Therefore, we had hypothesized that reward would facilitate representational convergence. The reason for this dissociation between the mPFC and anterior HC is unclear. The mPFC has been previously demonstrated to encode value-shaped representations of knowledge (Baram et al., 2021; Moneta et al., 2023; Paulus et al., 2021). Furthermore, both the mPFC and HC are linked to the brain’s reward circuit (Haber & Knutson, 2010), and the anterior portion of the HC is particularly connected to the mPFC (Adnan et al., 2016; Barnett et al., 2021; Poppenk et al., 2013). Specifically, post-encoding functional connectivity between the mPFC and anterior HC predicts subsequent memory transformation, both behaviorally (Audrain & McAndrews, 2022) and neurally (Tompary & Davachi, 2017). The anterior HC may also be particularly involved in motivational aspects of memory (Murty et al., 2017; Poppenk et al., 2013). Hence, one may have expected similar effects of reward on memory transformation in the anterior HC and mPFC.

However, consolidation is not complete after one night. Given that memory transformation may last for years, accompanied by a neural shift from HC to neocortex (Sekeres et al., 2018), it is possible that such reward effects on neural similarity emerge first in the HC and then shift to the mPFC at a later time point. Indeed, the HC may constitute a quick learning system that rapidly acquires not only episodic memory traces, but also regularities from similar events (Kumaran & McClelland, 2012; Schapiro et al., 2017). Through reward-biased replay, it may then coordinate the acquisition of generalized memory representations in the slower neocortical system (Kumaran et al., 2016). This may be one reason why, after a comparatively short time window of one day, we only observed reward effects on memory transformation in the anterior HC. Whether the differences between the mPFC and HC indeed reflect different time-courses of reward-enhanced memory transformation could be tested in future work using repeated retrieval sessions.

The hippocampus has been suggested to be functionally differentiated along its longitudinal axis, with more gist-like, schematic representations in the anterior HC, and more fine-grained, episodic representations in the posterior HC (Audrain & McAndrews, 2022; Guo & Yang, 2020; Poppenk et al., 2013). In particular, the (posterior) HC has been implicated in the reinstatement of low-level visual features (Bone & Buchsbaum, 2021), but also of events (Tompary & Davachi, 2017) and categories (Schultz, Sommer, et al., 2022). Our results only partially support this distinction. We indeed observed distinct result patterns along the longitudinal axis of the HC: The anterior HC showed increased memory transformation for high-reward retrieval trials. The posterior HC, on the other hand, did not yield evidence for episodic scene reinstatement.

Instead, we only observed scene reinstatement in the PHC. When participants correctly recalled a scene, the activation pattern in the PHC was more similar to the activation pattern during encoding of that scene. This is in line with previous research showing scene-specific pattern reinstatement in the PHC (Meyer & Benoit, 2022; Schultz et al., 2019; Schultz, Sommer, et al., 2022; Staresina et al., 2012) and scene-specific memory processing in general (Liang & Preston, 2017; Schultz et al., 2012; Schultz, Yoo, et al., 2022; Staresina et al., 2013). Anatomically, the PHC is a connecting hub between the dorsal visual stream and downstream regions in the MTL, including the entorhinal cortex and HC (Lavenex & Amaral, 2000; Suzuki & Amaral, 1994b, 1994a), and thus well-positioned to support spatial, scene-specific, or contextual memory (Eichenbaum et al., 2007).

The concurrent presence of original and transformed memory representations is consistent with accounts that the same memories exist in multiple forms at different levels of abstraction, with their relative strength of activation dependent on e.g. task demands (Gilboa & Moscovitch, 2021; Sekeres et al., 2018). Here, we observed that the same retrieval trials elicited both reinstated and transformed memory representations. These were present in the MTL and mPFC, respectively. Does this suggest that the two are independent of each other? Previous work has demonstrated that, during autobiographical retrieval, mPFC activity precedes and drives HC activity (McCormick et al., 2020, but see Campbell et al., 2018). Furthermore, a higher integrity of the anatomical connection between the HC and mPFC is associated with richer autobiographical memories (Williams et al., 2020). It is possible that the memory representations in the mPFC are instantiated earlier, but are by themselves insufficient to elicit successful episodic retrieval. However, if these are passed on through top-down modulation of the MTL (McCormick et al., 2020; Nawa & Ando, 2019; St Jacques et al., 2011), they may guide episodic retrieval and thus aid in recovering the details of a memory trace. Other methods with a higher temporal resolution, such as electrophysiological measures, may further elucidate these potential interactions between the complementary memory representations in the mPFC and MTL.

In summary, we have provided evidence for the concurrent activation of two types of memory representations – in the PHC of the original activity pattern that was present during encoding and in the mPFC and anterior HC of activity patterns that resemble transformed, generalized memories. Reward enhances neural memory transformation in the anterior HC, though it does not reliably promote episodic memory. Our results thus broaden our knowledge of the processes that lead to memory generalization, while motivating new questions about how reward shapes the structure of memory.

## Acknowledgments

We thank Matthias J. Gruber for sharing the experimental stimuli from Gruber et al. (2016), Steffen Jödecke for adapting the PHC masks and assisting with data collection, Ruud M.W.J. Berkers for generating the mPFC mask, and Nuno Busch, Martina Dietrich, Lena Kuschel, Adelina Kuzmanova, Mark Lauckner, Mewes Muhs, and Sarah-Lena Schäfer for assisting with data collection.

## Data and code availability

All data necessary to reproduce the reported results, i.e. behavioral data, ROI similarity values, and *t*-maps for the whole-brain searchlight analyses, are shared on OSF along with an R markdown file (https://osf.io/yracf/).

